# Coupling chromatin folding with histone modifications reveals dynamical asymmetry in the epigenetic landscape

**DOI:** 10.1101/2022.11.02.514881

**Authors:** Amogh Sood, Greg Schuette, Bin Zhang

## Abstract

Genomic regions adopt heritable epigenetic states with unique histone modifications, resulting in bistable gene expression without changes to the underlying DNA sequence. The significance of chromatin conformational dynamics to epigenetic stability is not well understood. We introduce a kinetic model to simulate the dynamic fluctuation of histone modifications. The model explicitly incorporates the impact of chemical modifications on chromatin stability as well as the contribution of chromatin contacts to the cooperativity of chemical reactions. Leveraging the model’s computational efficiency, we study the disparate time scales of chromatin relaxation and epigenetic spread to account for the recent discovery of both liquid and gel-like properties of chromatin. Strikingly different results were obtained for the steady state and kinetic behavior of histone modification patterns in fast and slow chromatin structural relaxation regimes. Our study suggests that the timescale of chromatin conformational dynamics maybe an important axis that biology fine tunes to regulate epigenetic stability.

---

Eukaryotic cells compactly package their genome into chromatin that consists primarily of nucleosomes formed by DNA wrapping around histone proteins [1, 2]. The core histones that make up the nucleosome are often subject to post-translational marking, including acetylation, methylation, phosphorylation, and ubiquitination [3, 4]. These marks partition chromosomes into distinct domains with differential transcription activity [5–9], providing active (euchromatic) and inactive (heterochromatic) regions with characteristic and non-overlapping chemical signals. The co-existence of chromatin stretches enriched and depleted of specific histone marks implies that epigenetic regulations exhibit multistability [9, 10].

Histone marks are known to promote (or restrict) the accessibility of the DNA sequence to transcriptional machinery [11–22]. By allowing a single DNA sequence to produce different gene expression patterns in different contexts, histone marks contribute to the emergence and maintenance of cell type diversity in multicellular organisms. Additionally, aberrant histone marks are frequently associated with cancer [23– 26]. Therefore, understanding the underlying mechanisms leading to the multistability of histone marks is pertinent to the study of a wide range of biological processes ranging from cellular differentiation to tumorigenesis [27–29].

Many theoretical models have been introduced to study the stability of histone marks [30–40]. Positive feedback underpins these models since existing marks will recruit enzymes to confer similar marks at new sites or nucleosomes [41–45]. This feedback relies on the 3D structure of the genome as chromatin loops facilitate the long-range spreading of histone marks by bringing nucleosomes far apart in sequence into spatial proximity [46–49]. On the other hand, chemical modifications could also affect nucleosome-nucleosome interactions, either by directly altering the physiochemical properties of amino acids or by recruiting additional protein molecules [50– 55], impacting chromatin organization. Therefore, chromatin is not an inert structure but rather an instructive scaffold and is inextricably linked to epigenetic regulation [56–58].

The importance of explicitly accounting for chromatin organization when studying histone marks is becoming increasingly appreciated in recent studies [59–68]. For example, many groups incorporate chromatin conformational dynamics by performing explicit polymer simulations when studying histone modifications [66–74]. Inherent to many studies is an assumption of fast chromatin structural relaxation compared to the timescale of chemical modifications. However, recent work reveals that chromatin exhibits very nontrivial rheology and viscoelastic properties, with multiple, disparate relaxation timescales, and organizes into regions of varying mobility [75–82]. In particular, *in vivo* studies of chromatin report solid-like behavior [83–85] and structural relaxation that occurs on the timescale of hours, a rate that is comparable to that of enzyme-mediated histone marks [39, 86, 87]. Therefore, the assumed timescale separation between chromatin structural and chemical dynamics in existing literature is not as clear and warrants further exploration.

Investigating the interdependence of chromatin structure and histone marks with more accurate time scale assumptions may help address an outstanding debate regarding the steady-state behavior of chromatin. Many existing models predict compact chromatin conformations regardless of the status of histone marks [59–68]. However, a more natural outcome would correspond to two states that support an open, unmarked (euchromatin) and collapsed, marked chromatin (heterochromatin). Designing models that support these two states would better represent biological systems and provide deeper insight into the stability of chromatin.

We present a theoretical model that explicitly accounts for the coupling between chromatin conformational dynamics and histone modifications (Figure 1). The model considers an array of **N** nucleosomes. The chemical state of chromatin at any given time, *t*, is denoted as a vector ***n***(*t*) ≡ {*n*_*i*_(*t*)} for *i* ∈ [1, **N**]. The binary variable, *n*_*i*_ ∈ {0, 1}, indicates the presence (or absence) of a histone mark at nucleosome *i*. We take inspiration from protein folding literature [88–91] and adopt a contact space representation of the chromatin conformation.

**FIG. 1.**
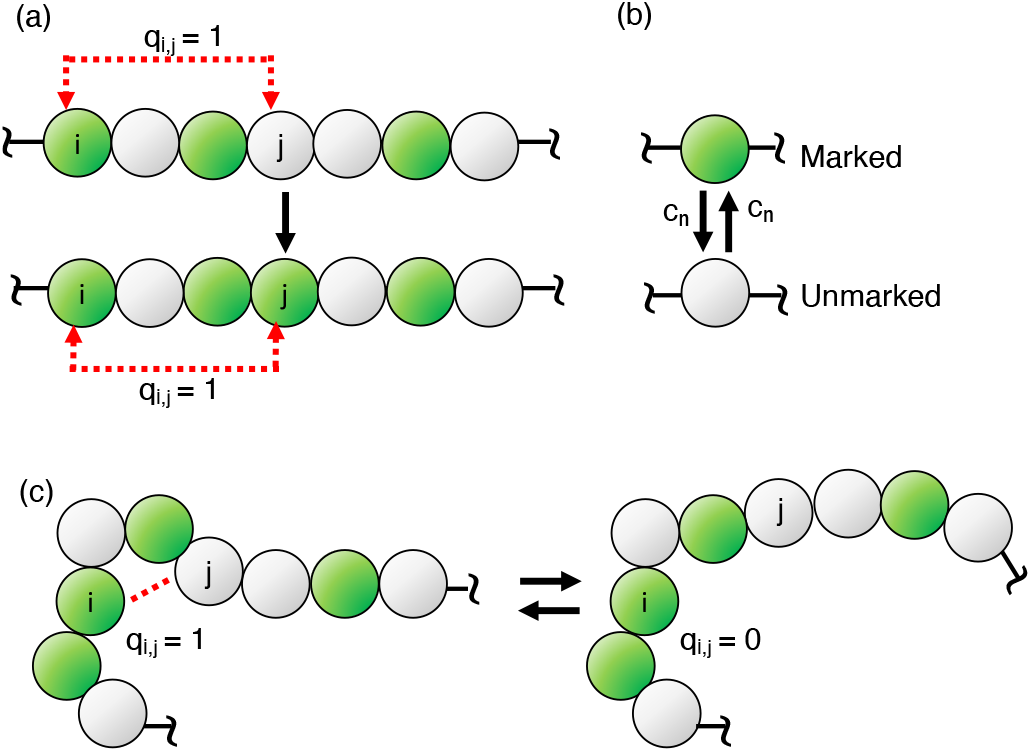
A schematic illustrating the salient features of the kinetic model that explicitly accounts for changes in histone marks and chromatin contacts and the coupling between the two. Nucleosomes are drawn as circles, with green and grey indicating marked and unmarked nucleosomes. (a) Marks can be added (or removed) in an enzyme-mediated recruited process wherein two sites that are in contact become similarly modified (Eq. 2). *q*_*i j*_ = 1 indicates that the two nucleosomes *i* and *j*, though far apart in a linear sequence, are in direct contact in the 3D space. (b) Nucleosomes can also be marked with on-site, random conversions that occur independent of chromatin contacts (Eq. 1). (c) Chromatin conformational dynamics is modeled as stochastic transitions in the contact space, where the contacts can form and break according to rates that depend on the polymer topology and nucleosome marks.

For example, we employ a vector of size **M, *q***(*t*) ≡ {*q*_*i j*_(*t*)} for *i, j* ∈ [1, **N**] and *j* − *i* > 1, to represent the conformation at time *t. q*_*i j*_ ∈ {0, 1} is again a binary variable that denotes the presence (or absence) of 3D contacts between a pair of nucleosomes (*i, j*). We assume that neighboring nucleosomes are always in contact, i.e., *q*_*i,i*+1_ ≡ 1.

Following previous studies [31, 33, 34], we consider two types of reactions that drive changes in histone marks. The first is an on-site, random conversion that arises from exchanging histone proteins with the nucleoplasm or from reactions catalyzed by non-cooperative enzymes. For example, the unmarked nucleosome *i* with *n*_*i*_ = 0, or 0_*i*_, can become marked with *n*_*i*_ = 1, or 1_*i*_, at a basal rate *c*_*n*_ that is independent of chromatin conformation and the state of other nucleosomes. Similarly, marked nucleosomes can be converted back to unmarked ones. The corresponding reaction schemes are denoted as

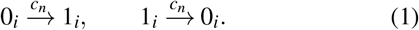

The second type of reaction represents recruited conversions, which, as a measure of cooperativity in the system, ensures that nucleosomes in spatial proximity are similarly marked. These reactions can arise due to the transfer of enzymes among nucleosomes in contact. We consider the cooperative effect for both addition and removal enzymes. Therefore, for a pair of contacting nucleosomes (*i, j*) in different chemical states, two outcomes can emerge as a result of recruited conversions. Either the mark at site *j* gets removed, or a new mark is introduced to site *i* as denoted below

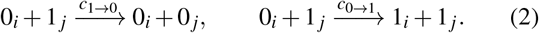

Both reactions occur with rate *c*_*r*_ unless otherwise specified.

Chromatin conformational dynamics are treated with analogous stochastic transitions. The rate of breaking and forming the contact between nucleosomes *i* and *j* is defined as *k*_*c*_ exp(− *βεn*_*i*_*n* _*j*_) and *k*_*c*_ exp(− *β*Δℋ), respectively. Here *k*_*c*_ is the basal rate constant and *β* = 1/*k*_*B*_*T* with *k*_*B*_ as the Boltzmann constant. *ε* = − 2.5*k*_*B*_*T* measures the interaction energy between marked nucleosomes, and is comparable to the value determined with a force spectrometer [92]. Δℋ = ℋ (***q***|*q*_*i j*_ = 1) − ℋ (***q***|*q*_*i j*_ = 0), where the free energy functional ℋ (***q***) is defined as

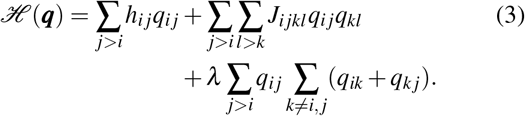

The linear term in the above expression, *h*_*i j*_, accounts for the entropic penalty of bringing nucleosomes *i* and *j* into contact [93]. The coupling term *J*_*i jkl*_ accounts for contact pair correlations, i.e. how forming contact (*i, j*) affects the probability of forming contact (*k, l*). These correlations are a result of polymer topology. The model parameters *h*_*i j*_ and *J*_*i jkl*_ were learned using a pseudolikelihood maximization approach [94]. The *λ* term in Eq. (3) accounts for the excluded volume effect by penalizing the formation of multiple contacts with the same nucleosome. Similar models have successfully been applied while studying the polymer dynamics of protein folding in contact space [90, 91]. Compared to polymer simulations in the Cartesian space, the binary contact model is computationally efficient and allows simulations across a wide range of timescales to interrogate the coupling between chromatin folding and histone marks. We note that simulation results are insensitive to the explicit form of the Hamiltonian defined in Eq. 3, and qualitatively similar results can be obtained from a mean-field expression (Fig. S1 and S2).

When chromatin conformational dynamics are not explicitly considered and collapsed chromatin structures are assumed to promote all contacts, the above model reduces to a mark-only version that has been studied extensively by many groups [33–35, 64, 66]. The mark-only model is known to exhibit bistability with two steady states where the fraction of marked nucleosomes is either close to 1 or close to 0 for large *c*_*r*_ values. We focus on a strongly cooperative regime wherein *c*_*r*_ = 100. Here and as follows, the values for all rate constants are reported in the unit of *c*_*n*_.

In line with contemporary literature, we first explore the regime where chromatin conformational dynamics are fast, and we chose *k*_*c*_ = 1000. Fig. 2a shows an example trajectory initialized with no chromatin contacts and completely unmarked nucleosomes. The blue and red traces depict the time evolution of the fraction of marked nucleosomes (⟨*n*⟩ ≡ Σ_*i*_ *n*_*i*_*/***N**) and fraction of contacts formed (⟨*q*⟩ ≡ Σ_*i j*_ *q*_*i j*_*/***M**), respectively. In the beginning, several transitions to the fully marked states are unsuccessful in the absence of sufficient contacts (Fig. 2b). However, as contacts build into the system, they endow the system with greater cooperativity and facilitate the spreading of marks through the recruited conversion pathway (Fig. 2a). Moreover, since marks confer attraction between sites, their establishment drives further collapse of the chromatin structure, and both contacts and marks increase in concert. Thus the two processes are intimately linked with each other, completing the transition to the formation of the marked and collapsed state. On the other hand, the coupling between chromatin folding and nucleosome modifications is less evident when the system transitions out of the fully marked state. As shown in Fig. 2c, a loss of marked nucleosomes proceeds without the vanishment of the chromatin contacts.

**FIG. 2.**
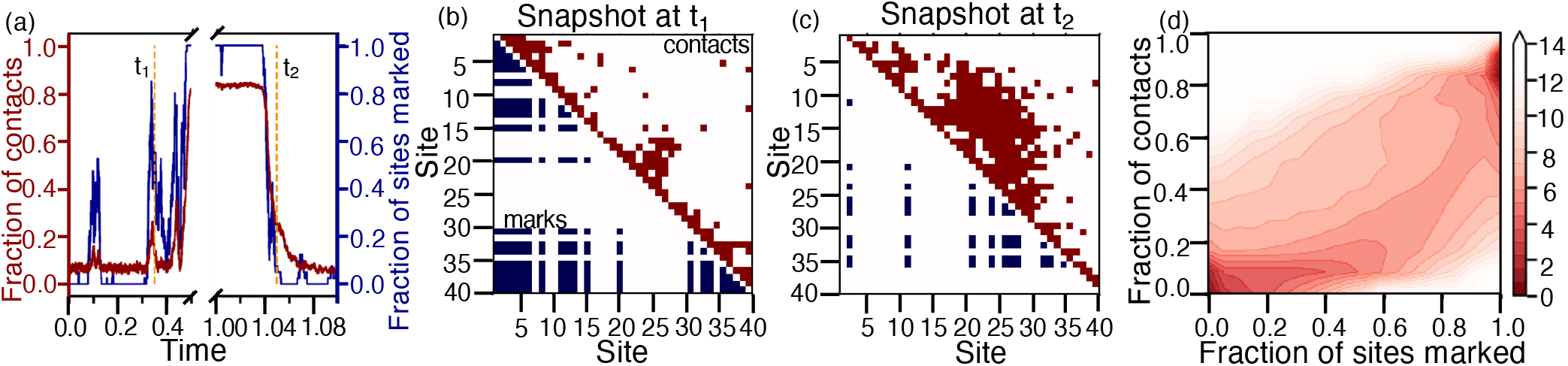
The model exhibits bistability with open, unmarked, and compact, marked chromatin in the regime with fast basal contact formation rate *k*_*c*_ = 1000. (a) Time evolution of the fraction of chromatin contacts (red) and the fraction of marked sites (blue) obtained from a simulation trajectory initialized from a state with zero histone marks and chromatin contacts. (b, c) Snapshots of the system showing contacts (upper triangle, red) and simultaneous marks (lower triangle, blue) between sites (*i, j*) taken at time *t*_1_ and *t*_2_ as indicated in part a. The two snapshots highlight an attempted transition to the marked state which fails due to insufficient long-range contacts in the system (b) and a transition from the marked state to the unmarked state leveraging recruited conversions via the existing long-range contacts (c). (d) Negative logarithm of the steady state distribution as a function of the fraction of marked sites and the fraction of chromatin contacts computed from 1.5 × 10^5^ *τ* long simulation trajectories. 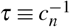.

To examine long-time behaviors of the model, we further computed the steady-state probability distributions as a function of ⟨*n*⟩ and ⟨*q*⟩, i.e., ℙ_ss_(⟨*n*⟩, ⟨*q*⟩). The negative logarithm of the distribution, which can be interpreted as a pseudopotential that quantifies the landscape of the stochastic system [86, 95–99], is shown in Fig. 2d. Two distinct steady states, a collapsed marked state and an open unmarked state resembling heterochromatin and euchromatin, respectively, are evident. The bistable behavior is consistent with the two-state switching kinetics shown in time evolution (Fig. 2a).

We performed additional simulations with slow chromatin conformation dynamics using *k*_*c*_ = 0.1 to explore the regime of retarded structural response expected in gel or solid materials. The time evolution of the system (Fig. 3a) is strikingly different from that shown in Fig. 2a despite being initialized with the same configuration. While the histone marks similarly transition between completely marked and unmarked states as in the fast chromatin case, chromatin contacts vary at a much slower rate. As a result, the intimate coupling between structure and sequence has disappeared. This is clear at the beginning of the trajectory, where the formation of even only a handful of non-backbone contacts seeds the spread of marks and supports cooperative transitions (Fig. 3b). Even though marks still confer attraction between sites in this system, they cannot drive a greater collapse of the polymer network because the chromatin relaxation rate is relatively slow and cannot respond before recruited conversions cause the epigenetic reaction network to transition back to the unmarked state. However, chromatin contacts do rise over time in response to the increase in the lifetime of the completely marked state, which was not robust at the beginning in the absence of sufficient contacts. The contacts tend to form in clusters and serve as nucleation sites for marks to enhance the cooperativity of epigenetic reactions. Similar to the lack of collapse seen in Fig. 3b, the contacts remain relatively unperturbed even when marks are lost since the contact dynamics are lethargic and cannot keep up with the marks as observed in Fig. 3c. After the initial equilibration, the dynamics occur on a network with relatively fixed connectivities. This is reflected in the steady-state behavior plotted in Fig. 3d, as the slow chromatin regime primarily exhibits a partially collapsed marked and a partially collapsed unmarked state.

**FIG. 3.**
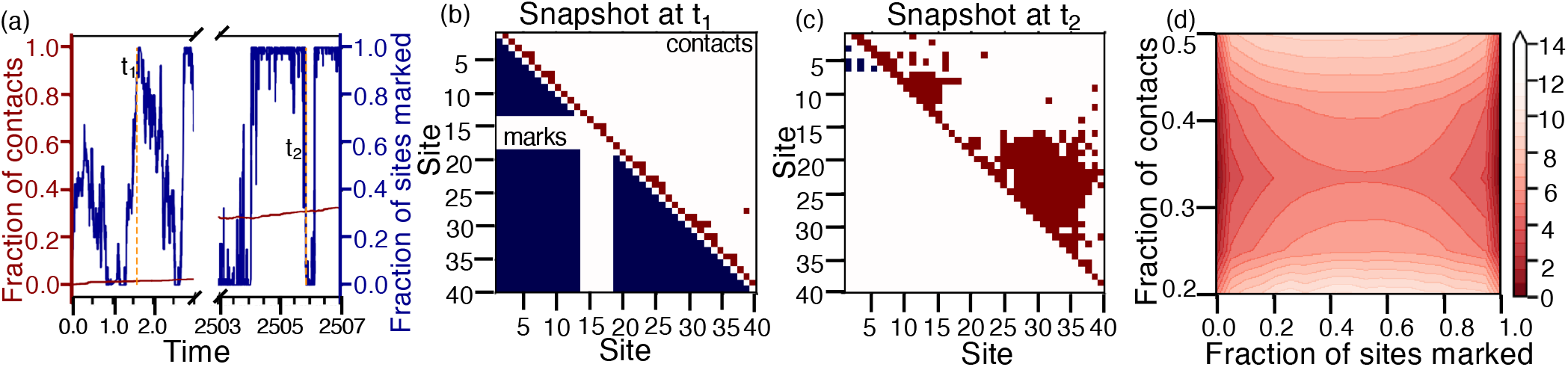
The model exhibits bistability with unmarked and marked chromatin in compact conformations in the regime with slow basal contact formation rate *k*_*c*_ = 0.1. (a) Time evolution of the fraction of chromatin contacts (red) and the fraction of marked sites (blue) obtained from a simulation trajectory initialized from a state with zero histone marks and chromatin contacts. (b, c) Snapshots of the system showing contacts (upper triangle, red) and simultaneous marks (lower triangle, blue) between sites (*i, j*) taken at time *t*_1_ and *t*_2_ as indicated in part a. The two snapshots highlight a transition to the marked state with only a handful of long-range contacts (b) and a transition between unmarked and marked states which occurs while the domains of long-range contacts remain relatively unperturbed (c). (d) Negative logarithm of the steady state distribution as a function of the fraction of marked sites and the fraction of chromatin contacts computed from 1.5 × 10^5^ *τ* long simulation trajectories. 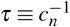.

While chromatin conformational distributions vary significantly in the fast and slow kinetics regimes, two steady states for marked and unmarked chromatin are observed in both cases. Fig. 4a shows the negative logarithm of the steady-state probability distributions as a function of the fraction of marked nucleosomes. For comparison, we also included the result from a model that neglects explicit dynamics of chromatin contacts by setting *q*_*i j*_ ≡ 1 for *i, j* ∈ [1, **N**]. As the reactions for histone marks are symmetric by design, the steady-state distribution is symmetric in the mark-only model. In the slow kinetics regime, the distribution remains symmetric because the contact dynamics are essentially decoupled from changes in histone marks. Since the fraction of contacts is less than one, the reactions for histone marks are less cooperative than the mark-only model, reducing the barrier for transitions between steady states. Strikingly, the distribution is asymmetric in the fast chromatin regime, with the marked state being the global minimum, and the entire landscape itself is tilted towards the unmarked state.

**FIG. 4.**
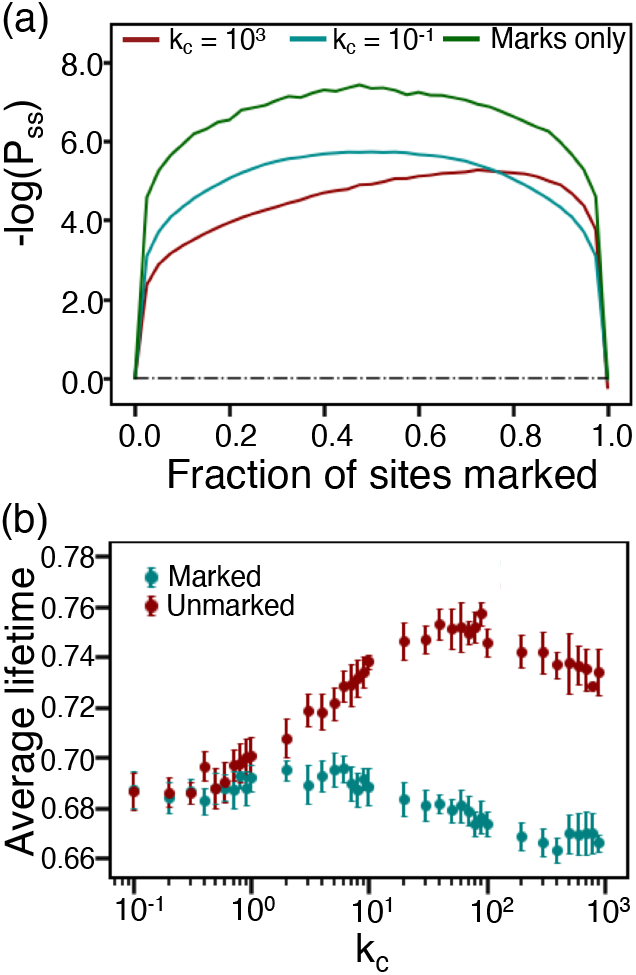
Coupling between histone marks and chromatin contacts introduces asymmetry in the epigenetic landscape and stabilizes euchromatin in the fast chromatin regime. (a) Negative logarithm of the steady state distribution for the fraction of marked nucleosomes computed with *k*_*c*_ = 1000 (red) and *k*_*c*_ = 0.1 (cyan). The result from a mark-only system that does not explicitly consider chromatin conformational dynamics with *q*_*i j*_ = 1 ∀ *i, j* is provided as a reference (green). (b) Variation in the average lifetime of marked (cyan) and unmarked states (red) with the basal chromatin contact rate *k*_*c*_.

**FIG. 5.**
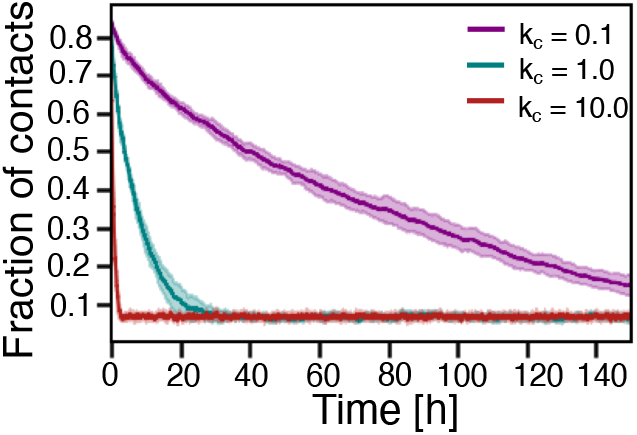
Resilience of collapsed chromatin conformations to perturbations in marks increases with decreasing *k*_*c*_. Plotted here is the change in the fraction of chromatin contacts as a function of time after removing cooperative reactions for modifying nucleosomes. The three curves correspond to simulations with basal chromosome contact formation rate *k*_*c*_ = 0.1 (purple), *k*_*c*_ = 1.0 (cyan), and *k*_*c*_ = 10.0 (red). When converting to the real-time unit, we used *c*_*n*_ = h^−1^ as estimated in Ref. 100.

To further understand the differences in the steady-state behaviors, we performed simulations at intermediate values of *k*_*c*_, the basal rate for chromatin structural dynamics. From these simulations, we determined the lifetime of both marked and unmarked states by partitioning the simulation trajectories into the two states [94]. As shown in Fig. 4b and S3, the lifetime of marked states remains largely unchanged by chromatin contact kinetics. On the other hand, the lifetime of unmarked states increases significantly as *k*_*c*_ is increased.

The imbalance between the lifetimes of steady states is consistent with the asymmetry in the landscape seen in Fig. 4a. It arises since in the collapsed state where most of the chromatin contacts are formed, the cooperative recruited conversions can drive transitions between marked and unmarked states with relative ease (Fig. 2c). However, in the open state, transitioning out of the unmarked state directly is harder due to a lack of contacts; the accumulation of chromatin contacts is slow due to the presence of entropic barriers inherent to the free energy functional ℋ (***q***) (Eq. 3). Spreading histone marks throughout the whole chain struggles to proceed with-out recruited conversions, leading to the frequently failed attempts seen in Fig. 2b. Overall, faster chromatin dynamics decrease the stability of heterochromatin relative to euchromatin. However, at extremely fast chromatin reorganization rates, chromatin contacts can reform substantially quickly in response to changes in histone marks. Therefore, the impact of diminished recruited conversions weakens, explaining the slight dip in the average lifetime of the unmarked state for large values of *k*_*c*_ in Fig. 4a. On the other hand, since the transitions between marked and unmarked states happen at rates much faster than the chromatin structural relaxation, the effect of contact kinetics on their relative lifetimes markedly disappears in the slow chromatin regime, with both marked and unmarked states having comparable average lifetimes.

Recent experiments paint a picture of chromatin as a highly heterogeneous system with variegated, non-trivial viscoelastic and rheological properties [75, 76]. Given these notions, we hypothesize that the timescale separation between the contact formation (breaking) in the chromatin network and the mark addition (removal) in the epigenetic reaction network can have profound consequences and is a potential modality cellular life can exploit. In particular, the strong coupling regime with fast chromatin kinetics might prove essential when silencing or activating gene expression during cell differentiation by reinforcing the effect of chemical modifications with accompanying structural changes in chromatin.

On the other hand, it is often important for biological systems to exhibit resilience against perturbations, and the slow chromatin regime can potentially endow the system with such resilience. For example, cells can leverage the slow chromatin regime to maintain a robust compact state when the histones become unmarked. To support this hypothesis, we carried out independent simulations with different chromatin conformational dynamics starting from a compact chromatin configuration with mostly marked nucleosomes. In contrast to the model used to obtain all the results presented so far, we removed recruited marking reactions by setting *c*_1→0_ = 100 and *c*_0→1_ = 0. This lack of cooperativity for spreading histone marks leads to a quick loss of marks in all cases on similar timescales regardless of chromatin dynamics (Fig. S4). However, the decompaction of chromatin, quantified by the decay of the fraction of contacts formed, showed dramatically different rates. For *k*_*c*_ = 0.1, the contacts can persist over hour timescales. A delayed response to changes in histone modifications is indeed consistent with observations from Ref. 101, where Eeftens et al. showed that chromatin remains compact upon a loss of condensates formed by enzymes that modify nucleosomes.

This work was supported by the National Institutes of Health (Grant No. R35GM133580).

