## Supplementary Material for "Coupling chromatin folding with histone modifications reveals dynamical asymmetry in the epigenetic landscape"

**CONTENTS**

I. Parameterizing the free energy functional of chromatin conformations in the contact space 2

Polymer conformations from molecular dynamics simulations 2

Parameter optimization with the pseudolikelihood approach 3

II. Details of Gillespie Stochastic Simulations 4

Estimating steady state lifetime from stochastic transitions 5

Estimating resilience of collapsed conformation to perturbations in marks 5

III. Mean field model 6

References 11

*

### I. PARAMETERIZING THE FREE ENERGY FUNCTIONAL OF CHROMATIN CONFORMATIONS IN THE CONTACT SPACE

For efficient simulations across a wide range of time scales, we consider chromatin conformational dynamics in the contact space. For a chromatin segment with **N** nucleosomes, the total set of contacts is represented with a vector of size **M**, $\boldsymbol{q}(t) \equiv \{q_{ij}(t)\}$ for $i, j \in [1, \mathbf{N}]$ and $j - i > 1$. The binary variables $q_{ij} \in \{0, 1\}$ denote the presence (or absence) of 3D contacts between a pair of nucleosomes $(i, j)$. We assume that neighboring nucleosomes are always in contact, i.e., $q_{i,i+1} \equiv 1$, and $\mathbf{M} = \mathbf{N}(\mathbf{N}-1)/2 - (\mathbf{N}-1)$.

We further define the free energy functional in the contact space as

$$\mathscr{H}(\boldsymbol{q}) = \sum_{ij} h_{ij} q_{ij} + \sum_{ijkl} J_{ijkl} q_{ij} q_{kl} + \lambda \sum_{ij} q_{ij} \Big( \sum_{t \neq i,j} q_{it} + \sum_{t \neq j,i} q_{tj} \Big). \tag{S1}$$

The linear terms $h_{ij}$ incorporate the entropic penalty of loop formation. We enforce translational symmetry such that contacts with the same sequence separation share identical penalty[1], i.e., $h_{ij} = h_{kl}$ whenever $j - i = l - k$. The correlation between contacts $(i, j)$ and $(k, l)$, which is caused by the polymer's topology, is accounted for by $J_{ijkl}$. Without loss of generality, the $J$ indices are ordered such that $k_1 \equiv j - i \leq l - k \equiv k_2$. Whenever $k_1 = k_2$, we additionally require $l > j$. Topologically-equivalent contact pairs can be identified by also defining $L \equiv l - j$ [2, 3]. Naturally, the topology-driven correlations are identical for topologically-equivalent contact pairs. Therefore, we assign identical $J$ values to all contact pairs with identical $k_1$, $k_2$, and $L$ values [2, 3]. Furthermore, we set $J_{ijkl} = 0$ for contacts with non-overlapping loops, which occur whenever $j \leq k$ or $l \leq i$, because the polymer's topology has no effect on these contacts' correlation [2, 3]. The third term accounts for the excluded volume effect that limits the number of contacts a given nucleosome can form.

We determined the parameters in Eq. S1 such that the free energy functional accurately describes the conformational distribution of polymers. As detailed below, the conformational distribution was produced with molecular dynamics simulations, and we used the pseudolikelihood approach for efficient parameter inference.

#### Polymer conformations from molecular dynamics simulations

We performed three independent molecular dynamics simulations to produce three conformational ensembles of polymers. The monomers are connected to nearest neighbors with the finite extensible nonlinear elastic (FENE) potential

$$u_{\text{bond}}(r_{i,i+1}) = -\frac{1}{2}KR_0^2 \ln\left[1-\left(\frac{r_{i,i+1}}{R_0}\right)^2\right], K_b = 30\varepsilon, R_0 = 1.5\sigma. \tag{S2}$$

The Lennard-Jones (LJ) potential was applied between all monomer pairs

$$U_{\text{wall}}(r) = \begin{cases} 4\varepsilon[(\frac{\sigma}{r})^{12} - (\frac{\sigma}{r})^6] + E_{\text{cut}}, & r < r_c \\ 0, & \text{otherwise,} \end{cases} \tag{S3}$$

where $E_{\text{cut}}$ is the energy of the LJ potential at the cutoff distance $r_c = 2.6\sigma$. We chose $\varepsilon$ as 0.3, 0.35, and 0.4 in the three simulations. The simulated polymer is 500 beads in length, and only the conformations of the central 40 beads were recorded. Simulating a longer polymer avoids potential edge effects that might produce different statistics for polymer beads at the boundary.

The LAMMPS software package [**?** ] was used to perform the simulations using reduced units and shrink-wrapping boundary conditions in reduced units with a time step of 0.005. The Langevin dynamics with a damping parameter of 10 were used to maintain the temperature. Temperature replica exchange was used with seven temperatures evenly spaced from 0.7 to 1.3, and data was collected from the replica with $T = 1.0$. Exchanges were performed every $100^{th}$ timestep, and configurations were collected once every 5000 timestep over one billion total timesteps. This yields $200,000$ configurations for each of the three homopolymers.

**Parameter optimization with the pseudolikelihood approach**

The model parameters $h_{ij}$ and $J_{ijkl}$ were learned using a pseudolikelihood maximization approach developed in previous work [4]. This approach adjusts the parameters to optimize the probability, or likelihood, of the conformations from a reference ensemble. From the 3D structures in Cartesian space obtained using molecular dynamics simulations, we obtained a list of contacts between monomer pairs using a distance cutoff of $1.707\sigma$. This conversion produces three ensembles of polymer conformations in the contact space, denoted as $\mathbb{B}_1$, $\mathbb{B}_2$, and $\mathbb{B}_3$, which were used in pseudolikelihood optimization.

The function used for parameter optimization is defined as

$$\begin{aligned}\ell_{\text{pseudo},\mathbb{B}}(\boldsymbol{h},\boldsymbol{J},\boldsymbol{\Delta h}) = &\sum_{b\in\mathbb{B}_1}\sum_{(i,j)}\log\left[\frac{1}{1+\exp\left(h_{ij}+\Delta h_1+\sum_{kl}J_{ijkl}q_{kl}^{(b)}\right)}\right]\\ &+\sum_{b\in\mathbb{B}_2}\sum_{(i,j)}\log\left[\frac{1}{1+\exp\left(h_{ij}+\Delta h_2+\sum_{kl}J_{ijkl}q_{kl}^{(b)}\right)}\right]\\ &+\sum_{b\in\mathbb{B}_3}\sum_{(i,j)}\log\left[\frac{1}{1+\exp\left(h_{ij}+\Delta h_3+\sum_{kl}J_{ijkl}q_{kl}^{(b)}\right)}\right]\end{aligned} \tag{S4}$$

Here, $\boldsymbol{h}$ ($\boldsymbol{J}$) is the model's full set of $h_{ij}$ ($J_{ijkl}$) parameters. The three terms on the right side correspond to the total pseudolikelihood of the energy function defined in Eq. S1 over the configurations from the three ensembles. Since the molecular dynamics simulations used to produce the three ensembles differ in the non-bonded interaction energy, we introduced $\boldsymbol{\Delta h} = \{\Delta h_1, \Delta h_2, \Delta h_3\}$ to account for the difference. $\Delta h_1$ was fixed at 0. Our use of three conformational ensembles with different degrees of polymer collapse provides a wide variety of conformations essential for probing the correlation between contact pairs.

Additional regularization was further introduced to ensure the robustness of parameter optimization, leading to the final objective function as

$$\ell_{\mathbb{B}}(\boldsymbol{h},\boldsymbol{J},\boldsymbol{\Delta h}) = \ell_{pseudo,\mathbb{B}} + \gamma\sum_{ijkl}J_{ijkl}^2 \tag{S5}$$

with $\gamma$ was set to 0.6. The function was optimized with the Broyden–Fletcher–Goldfarb–Shanno (L-BFGS-B) algorithm using SciPy version 1.5.2. All parameters were initialized at 0.

### II. DETAILS OF GILLESPIE STOCHASTIC SIMULATIONS

Stochastic simulations were carried out for the reaction network using an implementation of the Gillespie stochastic simulation algorithm [5] in Python using standard libraries. We set the parameters as follows: $\varepsilon = -2.5$, $\lambda = 0.01$, $c_n = 1.0\ \tau^{-1}$, $c_r/c_n = 100.0$. We simulate a system of size $\mathbf{N} = 40$ sites, and $\mathbf{M} = 741$ mutable, non-backbone contacts.

We ran simulations of length $3\times10^5\ \tau$ using different $k_c/c_n$ values, approximately corresponding to $\sim 10^8 - 10^{10}$ Gillespie moves. We discarded the first half of each trajectory to remove the influence of initial conditions on the simulation results. Steady-state probability distributions and contact maps were obtained by averaging over the remaining half of each simulated trajectory.

#### Estimating steady state lifetime from stochastic transitions

To better understand the origin of the asymmetry in the epigenetic landscape presented in Fig. 4 of the main text, we computed the steady-state lifetime as fellows. We identified the steady states using the fraction of modified nucleosomes as $\langle n \rangle \in (0.8, 1.0]$ and $\langle n \rangle \in [0.0, 0.2)$. To compute the average lifetimes of each, we sub-divided the second half of each of the simulated trajectories into $\sim 10$ sub-trajectories each of length $10^4 \tau$; this allows one to observe at least $\sim 10^2$ transitions between marked and unmarked states in each sub-trajectory. A successful transition from the unmarked to marked state was defined as the event wherein a system starting in the unmarked state arrives in the set defining the marked state (and vice versa). The lifetime of the unmarked (marked) state was defined to be the time between a transition to the unmarked (marked) state and transition to the marked (unmarked) state. The average lifetime was then obtained by computing the average time spent in each state in each sub-trajectory, and the error bars in Fig. 4 (b) reflect the standard error of the mean across the 10 sub trajectories.

We also tested the sensitivity of the results to the definition of marked (and unmarked) states used. We performed similar analysis by defining a marked (unmarked) state to be $\langle n \rangle \in (0.9, 1.0]$ ($\langle n \rangle \in [0.0, 0.1)$) and $\langle n \rangle \in (0.7, 1.0]$ ($\langle n \rangle \in [0.0, 0.3)$). Fig. S3 demonstrates qualitatively similar trends for the average lifetimes with varying $k_c$ using these thresholds to the ones seen in Fig. 4b of the main text.

#### Estimating resilience of collapsed conformation to perturbations in marks

We prepared a compact chromatin configuration with mostly marked nucleosomes. In contrast to the model used to obtain all the results presented so far, we removed recruited marking reactions by setting $c_{1\to 0} = 100$ and $c_{0\to 1} = 0$. The system was then allowed to relax at $k_c = 0.1, 1.0, 10.0$. For each value of $k_c$ we conducted 10 independent runs starting from identically prepared configurations. The solid color lines in Fig. 5 of the main text represents the average value computed across these 10 independent runs and the shaded regions represent the $\pm 1$ standard deviation.

The lack of cooperativity for spreading histone marks leads to a quick loss of marks in all cases on similar timescales regardless of chromatin dynamics as seen in Fig. S4

### III. MEAN FIELD MODEL

While the results presented in the main text were produced with the free energy functional defined in Eq. S1, the qualitative trends are rather robust to parameters in the functional. In fact, the qualitative trends can be reproduced by a mean-field free energy as a function of contacts ($\mathscr{F}(\langle q\rangle)$), provided in the absence of marks $\mathscr{F}(\langle q\rangle) \to 0$ as $\langle q\rangle \to 0$, and the addition of marks leads to more contacts building in the system and lends to $\mathscr{F}(\langle q\rangle)$ exhibiting two minima in the presence of attractive marks. A simple example would be to construct $\mathscr{F}(\langle q\rangle)$ as a quartic polynomial over $[0,1]$, where $\mathscr{F} = \sum_{r=0}^{4} a_r \langle q\rangle^r + \varepsilon_{ij} q_{ij}$, where $a_r$ are the polynomial coefficients of $\langle q\rangle^r$, and $\varepsilon_{ij}$ is a small attracting between two marked sites $(i,j)$.

To demonstrate the robustness of simulation results, we again performed stochastic simulations for the reaction network using an implementation of the Gillespie stochastic simulation algorithm [5] in Python using standard libraries. The rate for contact breaking and formation for a pair of nucleosomes $(i,j)$ was again defined as $k_c \exp\left(-\beta \varepsilon n_i n_j\right)$ and $k_c \exp(-\beta \Delta \mathscr{H})$, respectively. Now, $\mathscr{H}(\langle q\rangle)$ is defined as $\mathscr{H}(\langle q\rangle) \equiv \sum_{r=0}^{4} a_r \langle q\rangle^r + \log\left(\binom{\mathbf{M}}{\langle q\rangle \mathbf{M}}\right)$. The second term is needed in the microscopic model to account for the degeneracy in different configurations that yield the same $\langle q\rangle$ so that the macroscopic expression for $\mathscr{F}$ simplifies to a simple quartic polynomial. The parameters used were $a_0 = 0,\ a_1 = 150.0,\ a_2 = 664.043,\ a_3 = -6312.1,\ a_4 = 15000.0, \varepsilon = -0.55$. Simulations were carried out for $\mathbf{N} = 40$ ($\mathbf{M} = 741$) sites. For $k_c = 0.1, 1, 10$, we performed single simulations of length $2 \times 10^5\ \tau$. Three independent trajectories of length $5 \times 10^4\ \tau$ and $1.0 \times 10^4\ \tau$ were collected for $k_c = 100$ and 1000. The first half of each trajectory was again discarded to remove the influence of initial conditions on the simulation results. Steady-state probability distributions were obtained by averaging over the remaining half of each of the simulated trajectories.

A marked state was once again defined to be $\langle n\rangle \in (0.8, 1.0]$ and an unmarked state was defined to be $\langle n\rangle \in [0.0, 0.2)$. In order to compute the average lifetimes of each, we subdivided the remaining half of each of the simulated trajectories into $\sim 5$ sub-trajectories of equal length for $k_c = 0.1, 1, 10$. The average lifetime was then obtained by computing the average time spent in each state in each sub-trajectory, and the error bars in Fig. S1 reflect the standard error of the mean. For $k_c = 100, 1000$, the averaging procedure was done across the three independent trajectories. In our exploration, all of the main results could be recapitulated for this more minimal model and have been presented in Fig. S1.

In order to demonstrate that our results are robust to variations in system size, a larger system of size $\mathbf{N} = 60$ ($\mathbf{M} = 1711$) was also simulated using the mean-field model and parameters described above and a similar analysis was carried out (Fig. S2). Once again the qualitatively the features exhibited by the model remain unchanged.

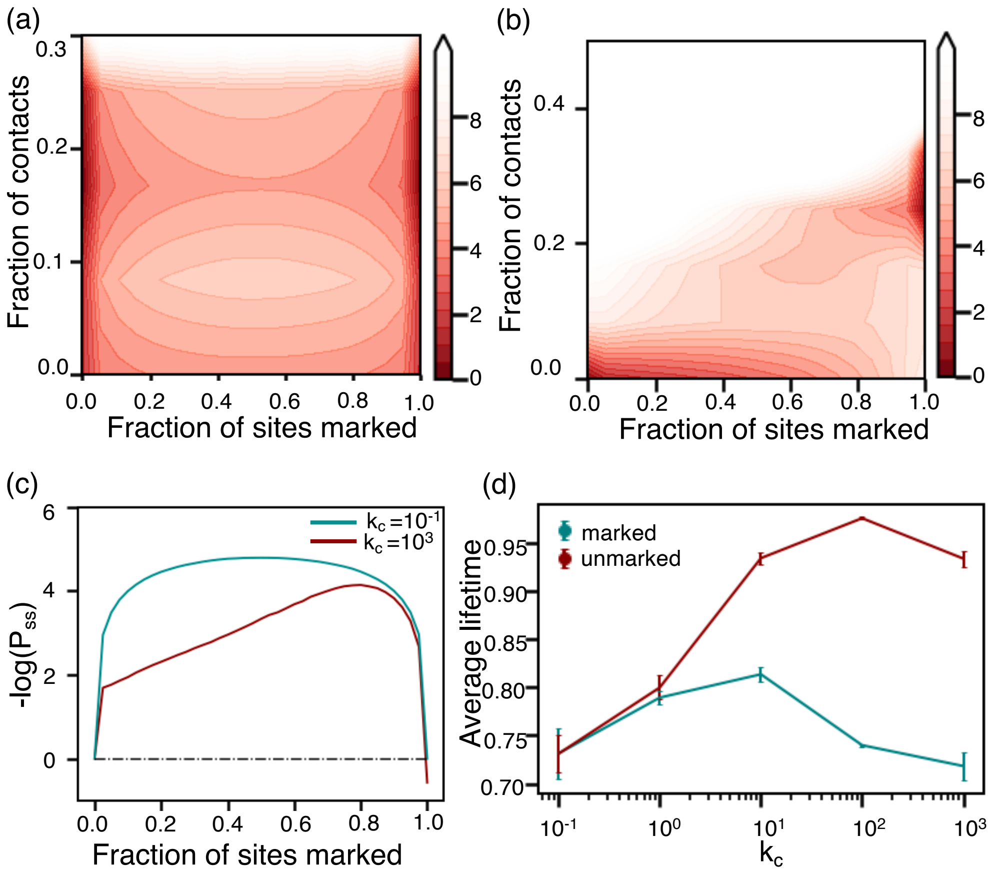


FIG. S1. A minimal mean field model recapitulates the main results presented in Fig 2, Fig. 3 and Fig 4 of the main text for a $\mathbf{N} = 40$ bead system. See text *Section: Mean Field Model* for additional discussions. Steady state probability distributions for the (a) slow chromatin regime with $k_c = 0.1$ and (b) fast chromatin regime with $k_c = 1000$. (c) $-\log(\mathbb{P}_{ss})$ for marks plotted as a function of the fraction of marked sites for $k_c = 1000$ (red) and $k_c = 0.1$ (cyan). (d) Variation in average lifetime of marked (cyan) and unmarked states (red) with $k_c$. See text *Section: Mean field model* for more details.

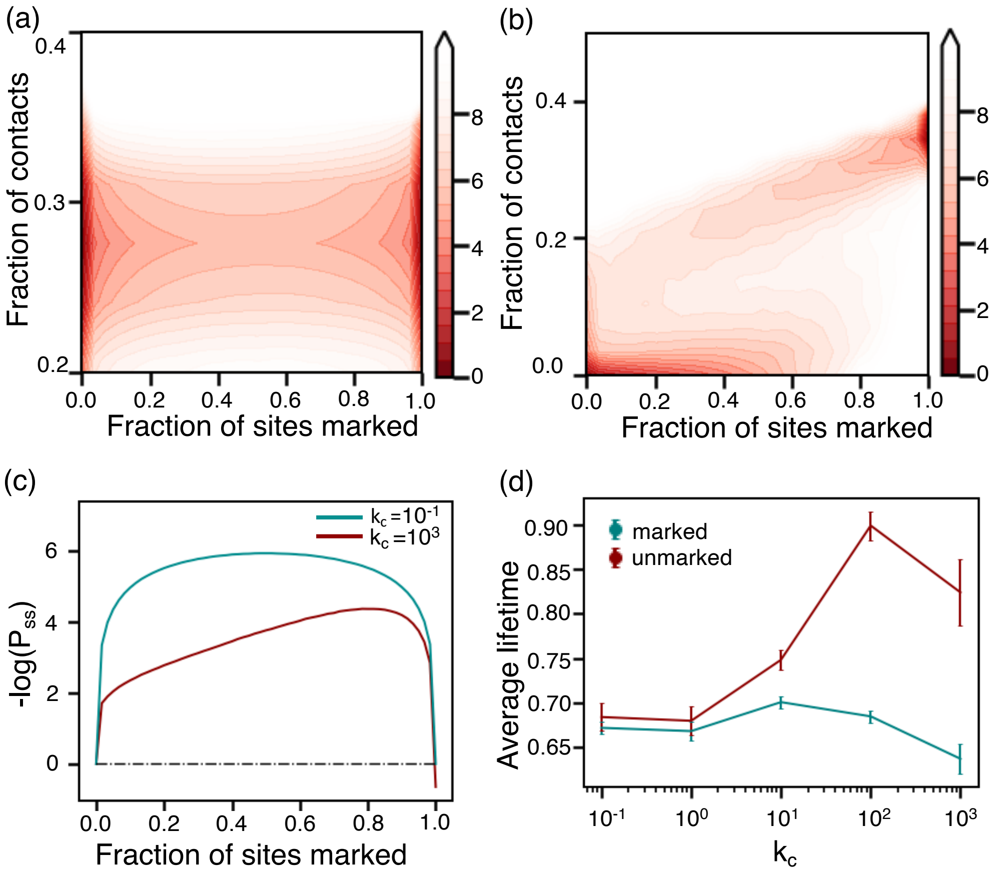


FIG. S2. A minimal mean field model recapitulates the main results presented in Fig 2, Fig. 3 and Fig 4 of the main text for a $\mathbf{N} = 60$ bead system. Steady state probability distributions for in the (a) slow chromatin regime with $k_c = 0.1$ and (b) fast chromatin regime with $k_c = 1000$. (c) $-\log(\mathbb{P}_{ss})$ for marks plotted as a function of marked sites for $k_c = 1000$ (red) and $k_c = 0.1$ (cyan). (d) Variation in average lifetime of marked (cyan) and unmarked states (red) with $k_c$. See text *Section: Mean field model* for more details.

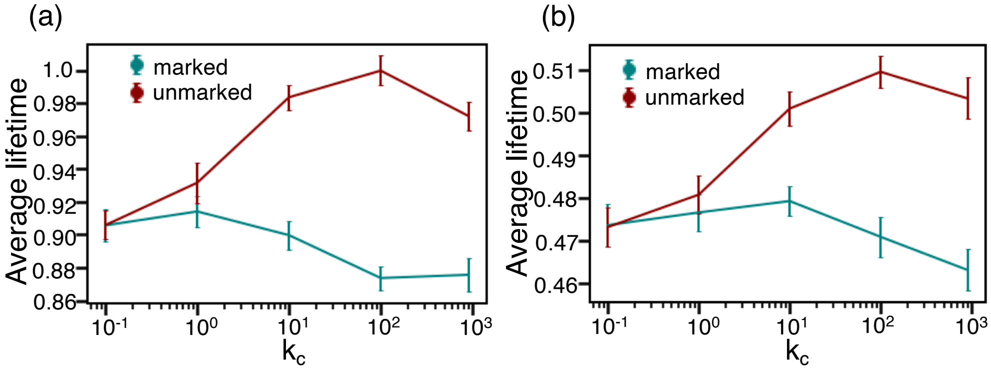


FIG. S3. Steady state lifetime estimation is robust with respect to the state definition. Qualitatively similar trends to Fig. 4b of the main text are observed for marked (unmarked) state defined as (a) $\langle n \rangle \in (0.9, 1.0]$ ($\langle n \rangle \in [0.0, 0.1)$) and (b) $\langle n \rangle \in (0.7, 1.0]$ ($\langle n \rangle \in [0.0, 0.3)$). See text *Section: Estimating steady state lifetime from stochastic transitions* for more details.

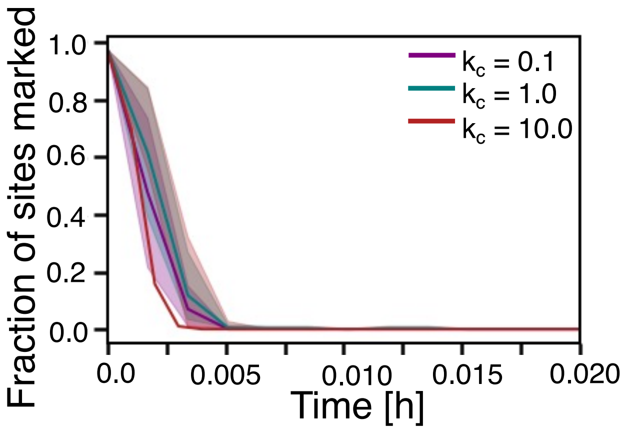


FIG. S4. Lack of cooperativity for spreading histone marks leads to a quick loss of marks in all cases on similar timescales regardless of $k_c$. Plotted here is the change in the fraction of sites marked as a function of time after removing cooperative reactions for modifying nucleosomes. The three curves correspond to simulations with basal chromosome contact formation rate $k_c = 0.1$ (purple), $k_c = 1.0$ (cyan), and $k_c = 10.0$ (red). When converting to the real-time unit, we used $c_n = 0.6\ \text{h}^{-1}$ as estimated in Ref. 6. See text *Section: Estimating resilience of collapsed conformation to perturbations in marks* for more details.
